# Comparative cell free salivary miRNA profile in *Bubalus bubalis* between diestrus and estrus stages

**DOI:** 10.1101/2024.09.08.611677

**Authors:** Rajeev Chandel, Dheer Singh, Suneel Kumar Onteru

**Author notes:** For Correspondence Dr. Suneel Kumar Onteru, M.V.Sc & Ph.D., <u>Email:</u>.

## Abstract

Despite buffaloes being primary farm animals, their reproductive performance remains poor mainly due to inaccurate estrus detection methods that ultimately has an economic impact on dairy industry as well as farmers. Recently, numerous studies showed potential of miRNAs as estrus biomarker. However, a miRNA profile of buffalo cell free saliva, a non-invasive fluid, at estrus and diestrus stages is missing. Hence, the present study was planned to identify differential levels of salivary cell free miRNAs in estrus as compared to the diestrus phase of buffalo oestrous cycle (n=3) in order to discover a possible estrus specific miRNAs as biomarkers. miRNA-Seq data analysis showed that in total 10 miRNAs i.e bta-miR-375, bta-miR-200c, bta-miR-30d, bta-let-7f, bta-miR-200a, bta-miR-12034, bta-let-7b, bta-miR-142-5p, bta-miR-2467-3p, bta-miR-30a-5p are significantly altered (log2foldchange >3 and p<0.05) during estrus in comparison to the diestrus phase in buffaloes, suggesting their estrus biomarker potential. Overall, 8 miRNAs i.e bta-miR-375 (6.87 Fold; p-value 0.003), bta-miR-200c (5.98 Fold; p-value 0.003), bta-miR-30d (4.17 Fold; p-value 0.015), bta-let-7f (3.34 Fold; p-value 0.022), bta-miR-200a (4.92 Fold; p-value 0.024), bta-miR-12034 (3.58 Fold; p-value 0.0025), bta-let-7b (3.06 Fold; p-value 0.031), bta-miR-30a-5p (4.7 Fold; p-value 0.036) were upregulated, whereas bta-miR-142-5p (-3.4 Fold; p-value 0.032) and bta-miR-2467-3p (-5.24 Fold; p-value 0.035) were downregulated during estrus. However, further validation study using qPCR is required in a large sample size in order to determine their estrus biomarker potential. In summary, our results revealed differential salivary cell free miRNAs profile during the oestrous cycle that may lead to the development of estrus specific miRNAs based point-of-care test applicable for the reproductive management of buffaloes in the field condition in the near future.

## 1. Introduction

Buffaloes (*Bubalus bubalis*) are primary dairy farm animal in South East Asia and the Mediterranean region [1]. Despite high milk and meat yield, their reproductive performance remains poor primarily due to silent estrus among various factors [1], [2]. Traditionally, estrus detection is done via visual observation of behavioral signs in an ineffective way. Even modern methods developed for estrus detection including heat mount detectors and activity monitors [3], [4], [5], [6] lack accuracy in buffaloes. As a result, half of the ovulations remains unnoticed [7], which causes a substantial financial loss to the dairy industry. Hence, accurate estrus detection method is urgently required to improve the conception rate and reproductive performance of buffaloes. Till now, numerous studies were performed to discover accurate estrus biomarker in various biofluids [8], [9], [10], [11], [12] from buffalo, but still a concrete field applicable biomarker is in search.

miRNAs are small non coding RNAs known to regulate their target protein levels either through mRNA degradation or preventing mRNA translation [13], [14]. Their potential as biomarker has been reported in earlier studies [15] owning to their association with various pathophysiological conditions in animals [16], [17]. In this context, circulating miRNAs in various biofluids including urine [18], saliva [9] and plasma [19] were reported as estrus biomarker in buffalo. Circulating miRNAs are the preferred biomolecules for biomarker discovery due to their better stability [20] and resistance to ribonucleases [21]**. T**hese circulating miRNAs may be differentially released into systemic circulation either actively or passively depending upon the physiological state of an animal by various organs including ovaries [22], [23], [24] and may ultimately appears in the saliva. Hence, it can be hypothesized that certain cell free salivary miRNAs may mirror the ovarian and plasma miRNAs expression [25], [26] with respect to the phase of an oestrous cycle. At last, circulating miRNAs can be detected in biofluids directly without RNA isolation using a simple, sensitive, and cheap LAMP [19] based method, which makes it feasible to use them as biomarker under field condition.

Saliva secreted by salivary glands is the most preferred biofluid for biomarker identification due to its ease of collection non-invasively [27], better stability [20], [21] and presence of different clinically useful molecules including nuclei acids [28]. Among different types of nuclei acids, miRNAs are frequently found in cell-free saliva [29]. Saliva being a partial ultra-filtrate of the plasma may contain certain bioanalytes of systemic origin [30], [31], [32] that can be used to determine pathophysiological state in an animal. In this context, it has been shown that bioanalyte profile of buffalo saliva differs during an oestrous cycle. For example, hormonal levels of estrogen and progesterone varies in buffalo saliva during oestrous cycle [33]. These studies enticed us to identify differential miRNAs levels in buffalo saliva during estrus and diestrus phase of the oestrous cycle.

In recent past years, miRNAs were discovered as estrus biomarkers in buffaloes on the basis of candidate miRNA approach. Unlike other biofluids, the characterization of total salivary miRNAs profile in buffaloes is yet to be done. Hence, the present study aimed to perform miRNA Seq on cell free total RNA isolated from buffalo saliva samples in order to identify candidate miRNAs with differential levels at the estrus and diestrus stage of an oestrous cycle that will be helpful to identify estrus specific biomarker.

## 2. Material and methods

### 2.1. Animals and estrus determination

Healthy Murrah buffaloes (n=3) were used as experimental animals after approval by the Institute Animal Ethics Committee. Animals were kept at the Livestock Research Centre, ICAR–National Dairy Research Institute, Karnal and maintained as per the standard conditions. Buffaloes in estrus were identified through visual observation of general behavioral signs such as sniffing/licking the vulva, flehmen’s reaction etc as well as gynaecoclinical examination for various parameters including cervical relaxation and vulva tumefaction and/or reddening of its mucous membrane. Estrus was further validated by the existence of dominant follicle (DF; ≥ 10mm) and lack of corpus luteum and the onset of ovulation through ultrasonography.

### 2.2. Saliva Collection and Processing

Saliva was aspired through 20 ml needless syringe from the buffalo lower jaw early morning before feeding and kept on ice. Saliva was diluted with PBS (1:1) and centrifuged at 3000g for 15 minutes at 4 °C to remove cells as well as the debris. 2 ml of diluted cell-free saliva (CFS) supernatant was added to and mixed well with 6 ml of TRIzol LS and incubated at room temperature for 5 minutes and then stored at −80°C till further use. Saliva samples were categorized as estrus (E) and diestrus (DE) stage on the basis of their collection on 0^th^ and 10^th^ day of an oestrous cycle, respectively.

### 2.3. Total RNA Isolation and quality assessment

Total RNA was extracted from CFS using TRIzol LS (Life Technologies, USA) as per the manufacturer’s protocol. In brief, the mix of CFS and TRIzol LS was thawed on ice and kept at room temperature for 5 min. 200ul chloroform was added per 1ml of Trizol LS used and the mixture was vortexed for 20 secs followed by incubation at room temperature for 5 minute and then centrifuged at 12000g for 15 minutes. Aqueous layer containing RNA was transferred to fresh Eppendorf tube and added 1ul of linear polyacrylamide, 0.1X of 3M NaOAc and 2.5X of absolute ethanol and kept at -80 O/N and finally centrifuged at 14000g for 30 minutes to get RNA pellet. Further processing of total RNA was performed as per manufacturer instructions and finally the extracted total RNA was DNaseI digested and purified using Zymo Columns, and final RNA elution was done in 10 μL of nuclease-free water and stored at −80°C. Total RNA was sent to Genotypic Pvt Ltd, Bangalore, India for custom NGS Library preparation and subsequent generation of miRNA Seq data. Total RNA purity and concentrations were determined using Nanodrop spectrophotometer, Qubit and Bioanalyzer available at Genotypic Pvt Ltd, Bangalore, India.

### 2.4. Small RNA library preparation and sequencing

Libraries for miRNA seq were prepared using QIAseq® miRNA Library Kit (Cat: 331502 Qiagen, Maryland, USA) as per following protocol. In brief, 5 ul of total RNA was used per sample to generate miRNA Seq library. Adapters were ligated sequentially to the 3’ (5’-AACTGTAGGCACCATCAAT-3’) and 5’ end (5’-GTTCAGAGTTCTACAGTCCGACGATC-3’) of miRNAs followed by their reverse transcription with primers containing Unique Molecular Index (UMI). cDNA prepared as mention above was further subjected to PCR amplification (22cycles) in order to enrich and barcode the reads in the cDNA library. Universal Illumina adapter sequence used in the protocol is 5’-AGATCGGAAGAGCACACGTCTGAACTCCAGTCAC-3’. Qubit fluorometer (Thermo Fisher Scientific, MA, USA) was used to quantify sequencing library. Agilent 2100 Bioanalyzer was used to analyze the fragment size distribution. The raw miRNA Seq data of Bubalus Bubalis cell free saliva samples were deposited in National Center for Biotechnology Information (NCBI) GEO database with accession no GSE275611. All libraries were sequenced through NovaSeq 6000 with 75 bp reads in order to generate greater than 10 million SE reads per sample as per manufacturer’s instructions.

### 2.5. In silico analysis of miRNA Seq data

miRNA Seq data was analyzed through various open-source softwares as described previously [34]. Reference mature miRNA file was downloaded from mirBase whereas genome fasta and GFF files were downloaded from NCBI. Reference indexes were created using bowtie build command. FastQC tool was used to assess the overall quality of each sample. UMIs were extracted from the reads and adaptors were removed using UMI_tools as mentioned previously [34]. UMI extracted and adaptor removed reads were filtered to retain reads for downstream miRNA analysis with read length >15 bp and <30 bp using Cutadapt tool. Filtered reads were aligned initially to known bta mature miRNA sequences from miRBase (v22) using Bowtie (with following parameters -n 1 -l 19 --norc -- best --strata -a). Unaligned reads from the previous step were aligned to the Bos Taurus reference genome GCF_002263795.3 using Bowtie (with parameters -n 1 -l 19 --norc -- best --strata -m 1). Samtools view option were used to sort and convert the SAM file into BAM file. BAM files were deduplicated using the UMI_tools dedup option –method=unique. Samtools idxstats were used to count each miRNA in every sample aligned to miRNAs whereas featurecounts tools were used to count reads aligned to miRNA coordinates in the reference genome using miRNA subset of GFF annotation file. Both mature miRNA-based counts and genome-based miRNA counts were combined into one file using R. DESeq2 was used to identify differentially expressed miRNAs and pheatmap function was used to create heatmap of log2 normalized counts.

### 2.6. miRNA target identification and functional analysis

The 3′UTR sequences for the Bos Taurus (GCF_002263795.3_ARS-UCD2.0_genomic) were downloaded from the UCSC Table Browser available at: http://genome.ucsc.edu. The sequences of 10 differentially altered miRNA identified in the present study were downloaded from miRbase v22.0 (http://microrna.sanger.ac.uk/sequences). miRanda tool [35] was used to identify miRNA target regions in the 3′UTR region of mRNAs using the following command: miranda mi.txt NCBI_RefSeq_3UTR -sc 140 -en -20 -out miranda_targets score where mi.txt is text file with miRNA Sequences, NCBI_RefSeq_3UTR file contains 3UTR of mRNAs, sc is score cutoff ≥□140, en is energy cutoff ≤□−20□kcal/mol. Functional identification of miRNA targets were determined through their enrichment in Biological process, Molecular function and Cellular component using Gene ontology database and KEGG pathway analysis using R. miRNet 2.0 [36] web based tool was used to perform network analysis of miRNA and its targets.

## 3. Results

### 3.1. RNA quantity and quality

Total RNA extraction has yielded sufficient quality of RNA for next-generation sequencing libraries preparation as shown in table 1. Nanodrop analysis of the total RNA showed concentration range of 2.2 to 39.6 ng/ul, with A260/A280 ratio from 1.49 to 2.34 and A260/A230 ratio from 0.11 to 0.46 indicating a good quality cell free RNA. Furthermore, Qubit analysis of total RNA showed concentration range from 2.48 to 4.4 ng/ul, indicating good cell free RNA yield. At last, percent of miRNAs in RNA isolated from Saliva samples ranges from 7 to 96% as per bioanalyzer analysis.

**Table 1.**
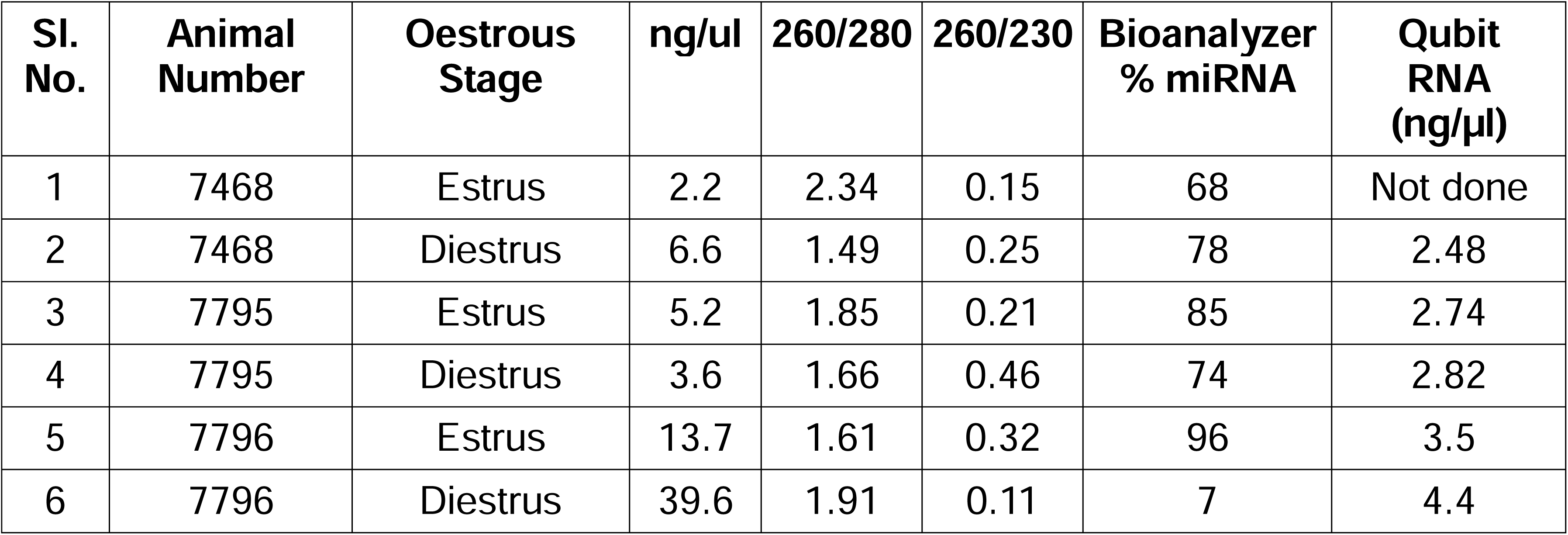
Total RNA concentration extracted from CFS samples.

| **Sl. No.** | **Animal Number** | **Oestrous Stage** | **ng/ul** | **260/280** | **260/230** | **Bioanalyzer % miRNA** | **Qubit RNA (ng/μl)** |
| --- | --- | --- | --- | --- | --- | --- | --- |
| 1 | 7468 | Estrus | 2.2 | 2.34 | 0.15 | 68 | Not done |
| 2 | 7468 | Diestrus | 6.6 | 1.49 | 0.25 | 78 | 2.48 |
| 3 | 7795 | Estrus | 5.2 | 1.85 | 0.21 | 85 | 2.74 |
| 4 | 7795 | Diestrus | 3.6 | 1.66 | 0.46 | 74 | 2.82 |
| 5 | 7796 | Estrus | 13.7 | 1.61 | 0.32 | 96 | 3.5 |
| 6 | 7796 | Diestrus | 39.6 | 1.91 | 0.11 | 7 | 4.4 |

### 3.2. miRNA Seq Libraries

To identify and characterize the miRNAs during the estrus and diestrus phases in buffalo saliva, miRNA Seq libraries were constructed from six individual CFS samples collected from 3 non-pregnant buffaloes on Day 0 and 10 of an oestrous cycle. The QIAseq miRNA Library kit used in the present study is specifically designed for generation of miRNA Seq libraries from low-input total RNA. The size of PCR amplified libraries is around the expected range for miRNA-sized library of approximately 180 bp as verified using TapeStation (Agilent) as shown in Fig 1(A-F).

**Fig. 1.**
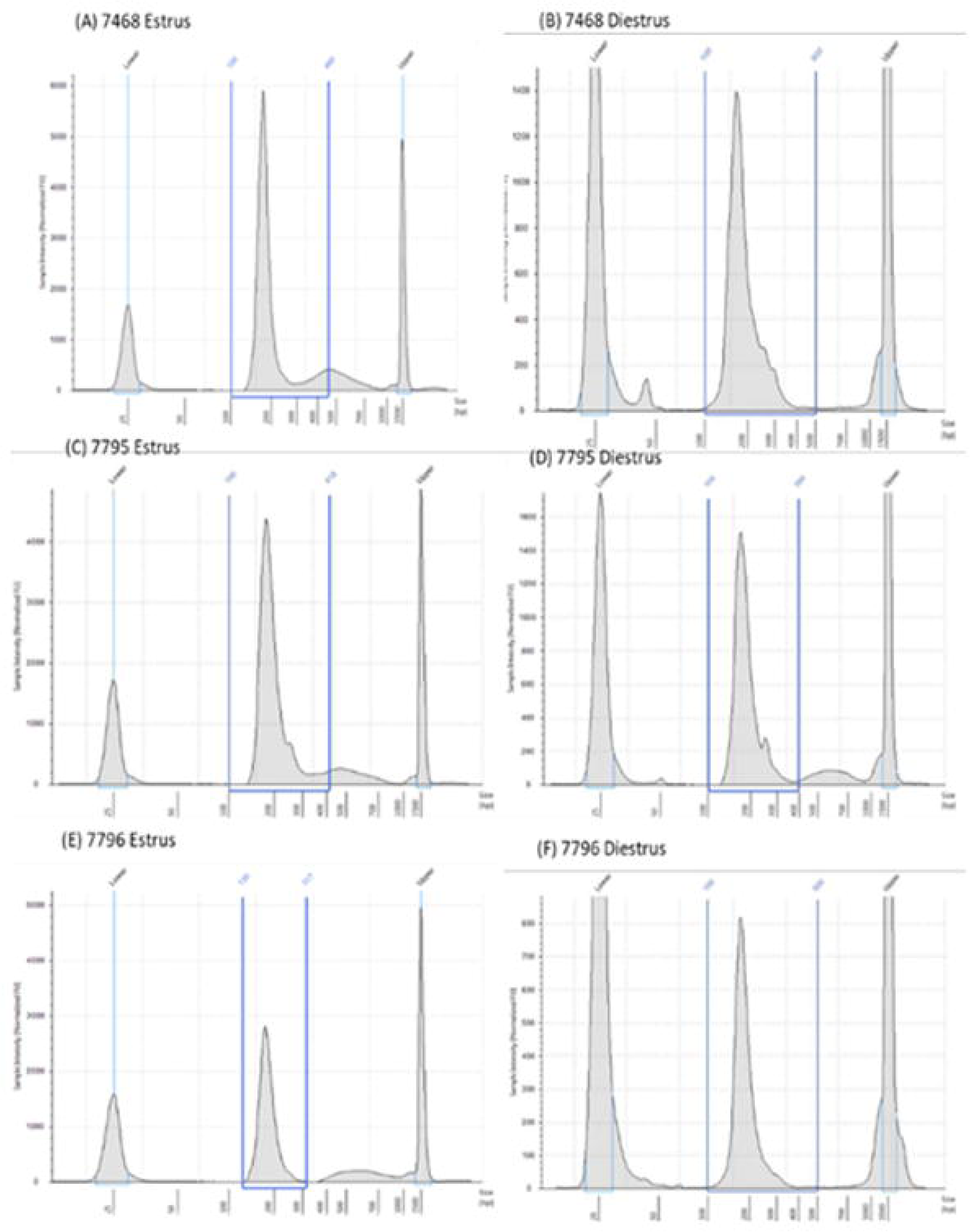
(A) to (F) miRNA Seq Libraries profiles as obtained on the TapeStation Instrument. First peak indicates lower Marker, second peak indicates cDNA, third pea’k indicates Upper Marker;

### 3.3. Quality Control of Sequencing Data

Six miRNA Seq libraries contained raw sequences in range from 10.7 to 34.8 million (Table 2). The miRNA-seq results revealed that the GC content ranges from 49 to 53%. In addition, the percentage of UMI extracted reads from the miRNA Seq libraries were between 20.46% and 97.52%. Filtering of UMI extracted reads for the read length 15-30 bp yielded 8.6 to 29.8 percent of raw reads. Finally, mapping of filtered reads to subset of mature bta-miRNAs from miRNA fasta file downloaded from mirBase v22 (Table 2) showed alignment rate from 0.04 to 0.23 percent during the first alignment step. Reads unaligned to mirbase were secondarily aligned to Bos taurus genome showed alignment rate from 1.98 to 3.12 percent. Our study identified overall 383 bovine miRNAs in buffalo saliva with 43 miRNAs being detected in all six samples used in the present study. On average, a total of 109 and 202 miRNAs with at least 1 read were detected in diestrus and estrus samples, respectively as shown in table 3.

**Table 2.** Summary showing various endpoints from miRNA sequencing of six saliva samples.

Reads preprocessing
| **Animal Number** | **Oestrous Phase** | **RAW READS** | **UMI EXTRACTED READS** | **FILTERED READS** | **mirBase aligned reads** | **Genome aligned reads** |
| --- | --- | --- | --- | --- | --- | --- |
| **7468** | Estrus | 34825999 | 7623081 | 3022666 | 32508 | 749665 |
| **7795** | Estrus | 16482345 | 3372610 | 1455185 | 9343 | 326034 |
| **7796** | Estrus | 16320984 | 15916109 | 4858305 | 38214 | 508605 |
| **7468** | Diestrus | 10668040 | 4034779 | 1187326 | 4137 | 248884 |
| **7795** | Diestrus | 12499884 | 3301763 | 1074926 | 7503 | 259976 |
| **7796** | Diestrus | 11463916 | 3846006 | 1677894 | 17030 | 348546 |

**Table 3.** Total miRNAs detected in six different CFS samples.

**Table No 2.** Summary showing various endpoints from miRNA sequencing of six saliva samples.
| **Animal Number** | **Oestrous Phase** | **Detected number of miRNAs** |
| --- | --- | --- |
| **7468** | **Estrus** | 207 |
| **7795** | **Estrus** | 128 |
| **7796** | **Estrus** | 271 |
| **7468** | **Diestrus** | 90 |
| **7795** | **Diestrus** | 119 |
| **7796** | **Diestrus** | 118 |

### 3.4. Differential gene expression analysis

DESeq2 was used for the differential expression analysis in saliva samples of estrus and diestrus groups and a total of ten miRNAs were found to be differentially expressed with Log2 fold change greater than 2.5 as shown in Volcano and MA plot (Fig 2a and 2b). Overall, 8 miRNAs i.e bta-miR-375 (6.87 Fold; p-value 0.003), bta-miR-200c (5.98 Fold; p-value 0.003), bta-miR-30d (4.17 Fold;p-value 0.015), bta-let-7f (3.34 Fold; p-value 0.022), bta-miR-200a (4.92 Fold; p-value 0.024), bta-miR-12034 (3.58 Fold; p-value 0.0025), bta-let-7b (3.06 Fold; p-value 0.031), bta-miR-30a-5p (4.7 Fold; p-value 0.036) were highly abundant, whereas bta-miR-142-5p (-3.4 Fold; p-value 0.032) and bta-miR-2467-3p (-5.24 Fold; p-value 0.035) were lowly abundant during the estrus as shown in Fig 3a and Table 4. Although, Principal component analysis (PCA) failed to show clear separation of estrus from diestrus phase samples (**Fig. 2c**), but PCA showed that estrus samples are spread out on PC1 axis as compared to diestrus samples which spread out more on PC2 axis suggesting that miRNAs explaining variability between samples of these two groups differs and hence their circulating levels may be useful for the identification of these two estrous phases respectively.

**Fig. 2.**
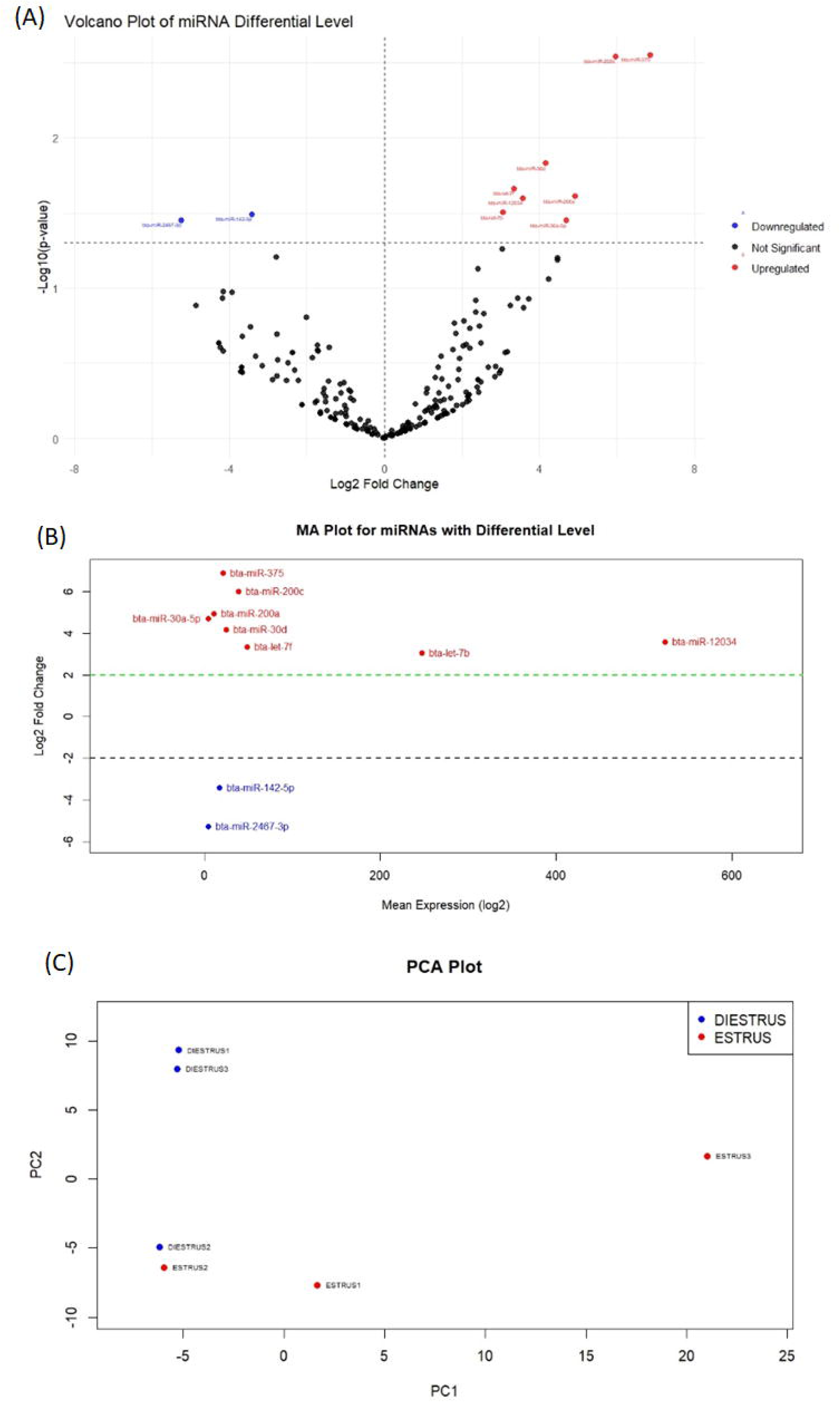
(A) Volcano plot showing fold-change of miRNAs versus statistical significance (P-values), in estrus relative to diestrus. MiRNAs with fold-change > 2 is marked by red vertical dotted lines and P < 0.05 is marked by red horizontal dotted line (B) MA Plot showing mean expression level of 1O differentially altered miRNAs (C) PCA plot showing separation of estrus and diestrus samples using the first two principal components.

**Fig. 3a.**
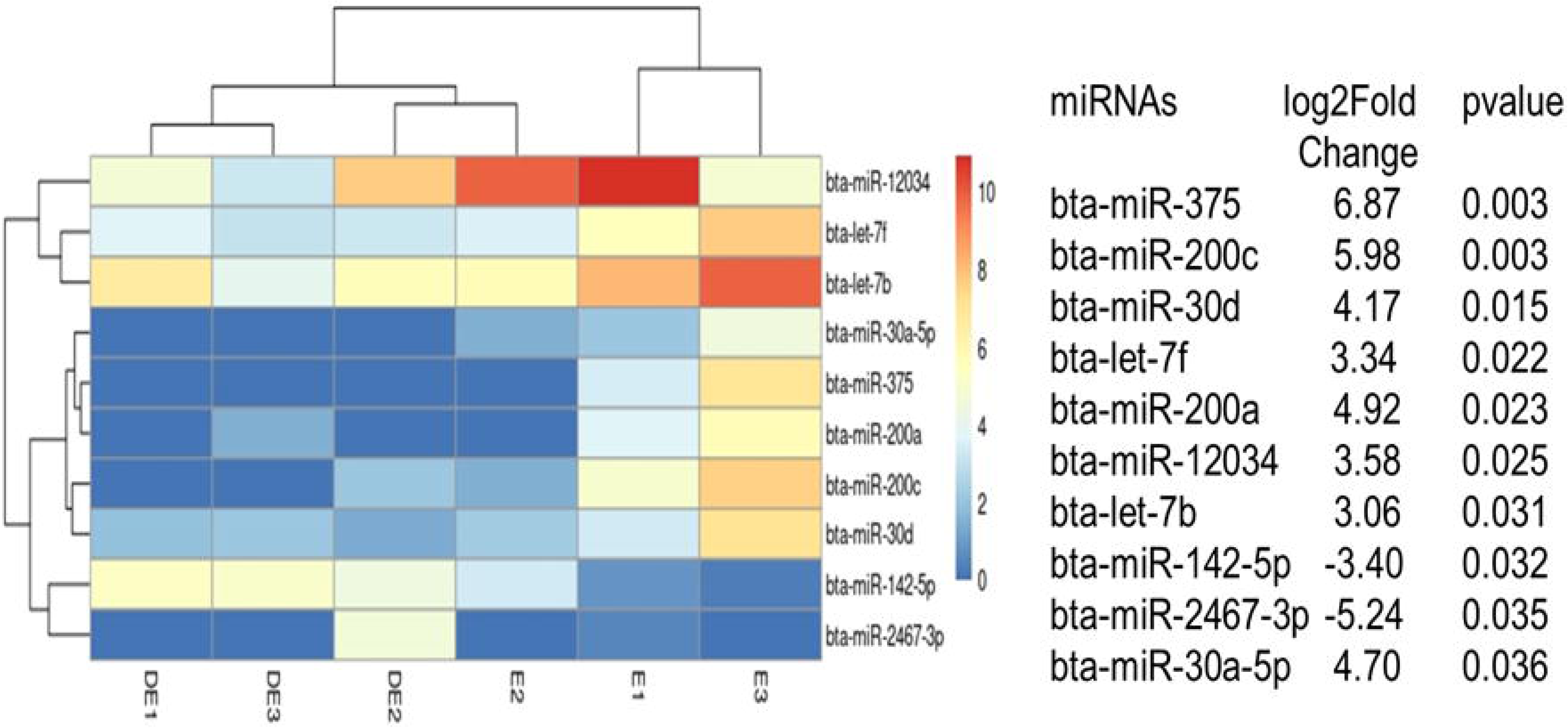
Heat map of top 10 miRNAs differentially expressed in estrus (n=3) and diestrus (n=3) samples of buffalo saliva where each row and column represent a miRNA and a sample respectively. The color scale illustrates the relative level of miRNAs. Table 4. Top 1O significant miRNAs which were highly or lowly abundant in buffalo saliva between the estrus and diestrus phases of the oestrous cycle as determined using miRNA sequencing.

**Fig. 4.**
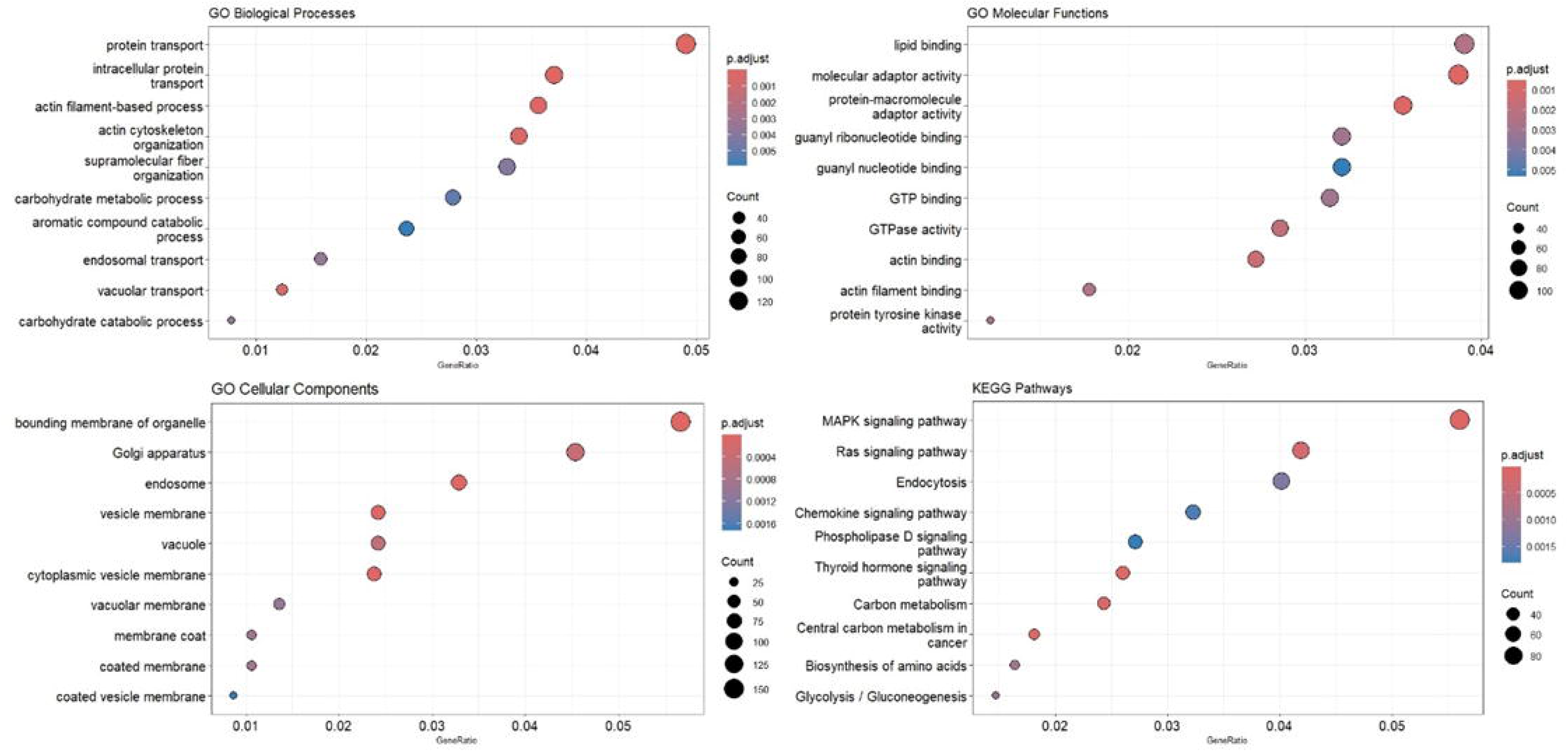
Gene Ontology (A-C) and KEGG (D) classifications of predicted targets regarding ten miRNAs with differential levels in E as compared to DE. CC cellular component, BP biological process, MF molecular function.

### 3.5. miRNA target prediction and analysis

miRanda 3.3a *(default version)* was used to predict miRNA targets using dynamic-programming alignment and thermodynamics. For *Bos taurus*, 7718 mRNAs sites are predicted to be targeted by 10 miRNAs. miRanda analysis showed that miRNAs identified in the present study regulates multiple target genes at the same time. For example, bta-miR-375 has 259 target sites, bta-miR-200c has 143 target sites, bta-miR-200a has 169 target sites, bta-miR-30a-5p has 164 target sites, bta-let-7f has 148 target sites, bta-miR-12034 has 1250 target sites, bta-let-7b has 812 target sites, bta-miR-142-5p has 7 target sites, bta-miR-2467-3p has 4622 target sites and bta-miR-30d has 14 target sites. miRanda analysis showed that among 10 miRNAs, bta-miR-2467-3p has highest number of targets sites i.e 4622 whereas bta-miR-142-5p has lowest number of targets sites i.e 7 in the 3’ UTR of mRNAs.

### 3.6. Functional and pathway enrichment analysis of miRNA targets

In order to elucidate the functional role of differential levels of salivary miRNAs in relation to oestrous cycle, 7718 mRNA targets as predicted using miRanda were used for gene ontology and KEGG pathway analysis in the present study. The predicted miRNA targets were broadly classified into biological processes, molecular functions and cellular components based on gene ontology (GO) (Fig 5. A, B and C). The top 5 biological processes enriched with predicted miRNA targets include protein transport (GO:0015031) 4.9%, intracellular protein transport (GO:0006886) 3.7%, actin filament-based process (GO:0030029) 3.56%, actin cytoskeleton organization (GO:0030036) 3.38%, vacuolar transport (GO:0007034) 1.23%. Further, top 5 molecular functions of predicted miRNA targets include protein-macromolecule adaptor activity (GO:0030674) 3.56%, molecular adaptor activity (GO:0060090) 3.87%, actin binding (GO:0003779) 2.72%, GTPase activity (GO:0003924) 2.86% and actin filament binding (GO:0051015) 1.78%, among others. At last, top 5 cellular components pathways enriched with predicted miRNA targets are bounding membrane of organelle (GO:0098588) 5.67%, Endosome (GO:0005768) 3.29%, Vesicle membrane (GO:0012506) 2.42%, Cytoplasmic vesicle membrane (GO:0030659) 2.38% and Golgi apparatus (GO:0005794) 4.54%. Top 5 KEGG pathways enriched with miRNA targets are MAPK signaling pathway (bta04010) 5.6%, Central carbon metabolism in cancer (bta05230) 1.81%, Thyroid hormone signaling pathway (bta04919) 2.6%, Carbon metabolism (bta01200) 2.43% and Ras signaling pathway (bta04014) 4.19%.

**Fig. 5:**
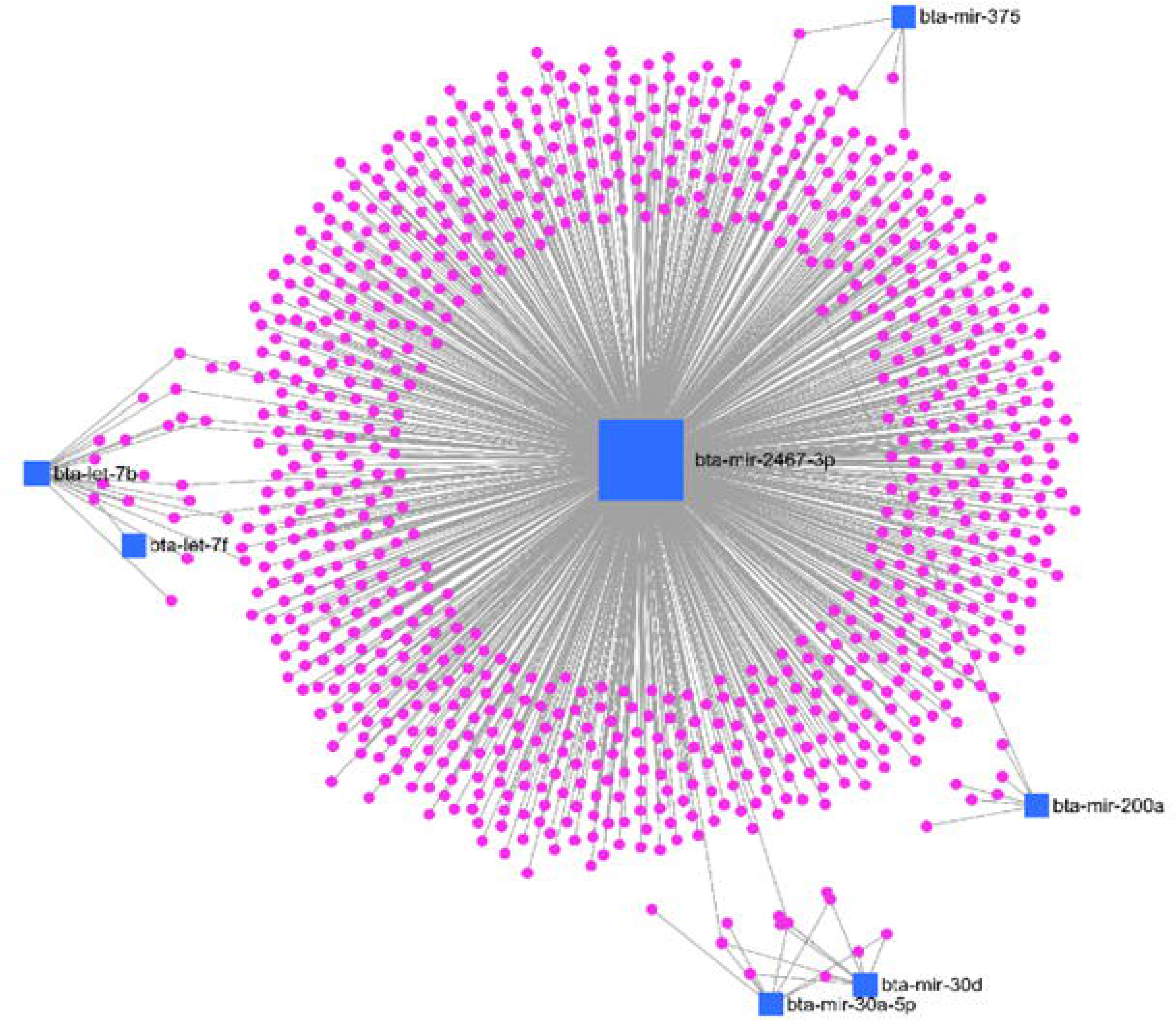
miRNet 2.0 Network analysis: showing miRNA regulating different genes. Genes are highlighted as pink circles, whereas miRNAs are represented as blue squares.

### 3.7. Network analysis of miRNA and its targets

miRNet 2.0 analysis of differentially present miRNA revealed miR-2467-3p is having highest degree centrality followed by bta-let-7b. It suggests that miR-2467-3p is regulating a greater number of genes that may be involved in pathways related to oestrous cycle regulation (Figure 5).

## 4. Discussion

The present study was planned to identify estrus specific salivary miRNA as biomarker in buffalo. Saliva was preferred over other biofluids for biomarker discovery due to its ease of collection non-invasively from farm animals. Total RNA isolated from cell free saliva sufficiently produced good RNA yield. This RNA yield is adequate enough to generate miRNA-Seq libraries using Qia Seq miRNA Library kit. Despite successful generation of miRNA-Seq libraries, there are several bottlenecks to overcome in the future before the use of miRNAs as biomarkers including optimization of RNA isolation method for extraction of high quantity RNA [37], [38], [39], [40], [41]. In addition, saliva contains abundant microbial species that needs to be addressed as compared to various other biofluids which are usually free of microbes [42].

To discover estrus-specific miRNAs, small RNA sequencing is considered the gold standard method mainly due to its high throughput capability and its ability to detect and quantify isomiRs as well as novel miRNAs [43], [44]. In this study, 0.35-1.08% of filtered and clean miRNA reads were aligned to Bos taurus miRNAs annotated in miRbase suggesting a large portion of the original reads are exogenous small RNAs derived either from bacteria or from dietary plants. In concordance, it has been reported previously that RNA-Seq reads generated from human cell free saliva RNA aligns approximately 8-17% to the human genome [45].

Our study outlines the possibility of using cell-free salivary miRNAs as an estrus biomarker by identifying a subset of ten cell free salivary miRNAs with higher levels of bta-miR-375, bta-miR-200c, bta-miR-30d, bta-let-7f, bta-miR-200a, bta-miR-12034, bta-let-7b, bta-miR-30a-5p and lower level of bta-miR-142-5p and bta-miR-2467-3p at estrus as compared to the diestrus phase in buffalo. Among these identified miRNAs, expression and role of bta-miR-12034, bta-miR-30a-5p and bta-miR-2467-3p is yet to be determined in the context of estrous cycle and reproduction of bovine.

One of the studies reported the expression of bta-miR-375 in granulosa cells of preovulatory dominant follicles [46] suggesting its role in follicular growth and oocyte maturation via regulation of transforming growth factor beta signaling pathway [47]. Higher salivary levels of miR-375 in buffalo observed in the present study is in concordance to the previous study reporting its higher levels during estrus in cows [48]. Moreover, earlier studies reported extracellular presence of miR-375 in exosomes [47] and microvesicles [49] suggesting its possible presence in vesicular fraction of buffalo saliva.

Higher salivary levels of let-7b observed during estrus in the present study is supported by the previous study reporting lower let-7b levels in early and progressed atretic follicles in comparison to healthy follicles suggesting its role in cell survival [50], [51]. It is one of the most abundant miRNAs in mural GCs and CCs [52] and has a pro-angiogenic role in corpus luteum and ovulation as well as in the regulation of FSH secretion [53], [54], [55]. MiR-200a is found to be highly expressed in ovaries [56] suggesting its role in ovarian development [57], [58] probably through its migratory, proliferative, and invasive activity [59], [60], [61]. Furthermore, miR-200a and miR-200c have a role in ovarian follicular and luteal development [53], [58], [62] and their downregulation ends the anestrous phase in sheep [63]. miR-200a/b/c may play a key role in estrus during the non-breeding season [64]. In contrast to our findings, Singh et al 2016 did not observe any change in miR-200c levels in buffalo saliva during oestrous cycle [25], suggesting that the lack of agreement might be due to differences in RNA isolation protocol and/or analytical platform used.

Higher salivary levels of bta-let-7f observed in the present study is in agreement to earlier study reporting higher plasma levels of bta-let-7f during estrus in cows [26]. Furthermore, earlier studies showed higher expression of bta-let-7f in ovary [65], [66] during proestrus and estrus phase of an oestrous cycle [67], suggesting its release from ovary into systemic circulation, although it is also expressed in various other tissues [68], [69], apart from ovary. It has a role in the follicle-to-luteal transition [46], [70], [71]. The bta-miR-30d is also highly expressed in ovaries of multiple species [65], [66], suggesting its housekeeping role in ovarian physiology [56] and in the follicle-to-luteal transition [46], [71].

In support of lower levels of salivary bta-miR-142-5p during estrus in the present study, it has been reported that bta-miR-142-5p is highly expressed in granulosa cells of dominant follicle at day 7 as compared to day 3 of the oestrous cycle [71] and has a role in granulosa cell survival via regulation of the Slit/Robo pathway activity [72]. Another study reported reduced expression of miR-142-3p in follicular tissues in comparison to the luteal tissues of ruminants [23], [70] further supporting our data.

In order to better understand the physiological functions and biological processes regulated by miRNAs identified in the present study to be at the significantly differential levels in buffalo saliva during estrus and diestrus phase, miRNA target genes were predicted using miRanda tool. Overall, pathway analysis of 7718 predicted targets of ten differentially altered miRNAs identified in the present study revealed that MAPK signaling pathway, Central carbon metabolism in cancer, Thyroid hormone signaling pathway, Carbon metabolism and Ras signaling pathway are the top five affected KEGG pathways. Among these pathways, p38 MAPKs [73], [74], [75] and Ras signaling [76] and thyroid hormones [77] pathways were reported to be involved in various aspects of the mammalian estrous cycles. Although, functional analysis of differentially altered miRNAs and their target genes suggested their role in estrous cycle, but the origin of these miRNAs needs to be verified before their use as estrus biomarker.

Our data differs from the previous studies due to various factors that influences miRNA levels and profile in biofluids including the differences in protocol for RNA isolation, miRNA quantification and normalization, sequencing libraries preparation [78], [79], [80] and bioinformatics pipeline used. In addition, lack of agreement between studies might be due to variability in the reproductive physiology of experimental animals used. The present study showed presence of very few cell free miRNAs in buffalo saliva that too with low levels, suggesting that abundantly expressed isoform of each miRNA in specific biofluid may differ from those listed on miRBase.

Our study has a number of limitations. First, the differential miRNA profile generated in the present study is from relatively small number of saliva samples. Hence, the results of the present study must be validated in large sample size using qPCR. Second, our findings must be validated in different buffaloes’ breeds. The third limitation is an inability to specifically pinpoint the origin of cell free salivary miRNAs in order to determine their implication during the oestrus cycle. Fourth, miRNA Seq library preparation kit from Qiagen used in the present study is susceptible to adaptor ligation bias that may lead to erroneous quantification of miRNA [81]. However, the cell free miRNA profile observed in biofluid depends upon the processing methods such as extraction of RNA and processing procedures. Furthermore, erroneous quantification of reads is caused by PCR amplification of RNA Seq libraries. In the present study, reads deduplication was done to remove PCR bias through unique molecular identifiers (UMIs) based information, but their effectiveness in small RNA-Seq data analysis is unclear [82], [83]. This biased effect may be higher in biofluids due to lower miRNA content [84] thereby necessitates caution while arriving at any meaningful conclusion and warrants validation of our findings in the future.

In future, use of salivary miRNAs as estrus biomarker in the field condition requires establishment of guidelines and protocols for saliva collection, processing and storage in order to preserve the salivary RNA profile as close to the original as possible via minimizing the effect of the preanalytical variables on RNA quality and quantity [85], [86]. In addition, one must take into account variability in salivary cell free miRNAs profile from the cellular miRNA after lysis and inaccurate volume measurement of the viscous saliva [87]. At last, method needs to be developed that can detect cell free miRNA in saliva, preferably directly without RNA isolation, through RT-LAMP is a promising tool for this purpose [19].

## Conclusion

In summary, our study for the first-time yielded insights of salivary miRNAs profile in buffalo using miRNA Seq technology and identified ten salivary cell-free miRNAs with estrus biomarker potential. This study expands our understanding of cell free salivary miRNA profile in buffalo during oestrous cycle. Hence, the output of the study may be useful in the future to detect estrus specific single or a panel of miRNAs using an easy, reliable and cheap field applicable method.

## Acknowledgments

The authors thank the Director, ICAR-National Dairy Research Institute, Karnal, Haryana, for facilitating the work, and Bill and Melinda Gates Foundation (Grant number: OPP1154401) for financial assistance.

## References

[1] S. Mondal, B. S. Prakash, and P. Palta, “Endocrine Aspects of Oestrous Cycle in Buffaloes (Bubalus bubalis): An Overview,” Asian-Australas J Anim Sci, vol. 20, no. 1, pp. 124–131, Nov. 2006, doi: 10.5713/AJAS.2007.124.

[2] V. S. Suthar and A. J. Dhami, “Estrus detection methods in buffalo,” 2010.

[3] K. K. Verma et al., “Characterization of physico-chemical properties of cervical mucus in relation to parity and conception rate in Murrah buffaloes,” Vet World, vol. 7, no. 7, pp. 467–471, 2014, doi: 10.14202/VETWORLD.2014.467-471.

[4] R. M. Selvam, S. K. Onteru, V. Nayan, M. Sivakumar, D. Singh, and G. Archunan, “Exploration of Luteinizing hormone in murrah buffalo (Bubalus bubalis) urine: Extended surge window opens door for estrus prediction,” Gen Comp Endocrinol, vol. 251, pp. 121– 126, Sep. 2017, doi: 10.1016/J.YGCEN.2016.12.002.

[5] J. Roelofs, F. López-Gatius, R. H. F. Hunter, F. J. C. M. van Eerdenburg, and C. Hanzen, “When is a cow in estrus? Clinical and practical aspects,” Theriogenology, vol. 74, no. 3, pp. 327–344, Aug. 2010, doi: 10.1016/J.THERIOGENOLOGY.2010.02.016.

[6] R. M. Selvam and G. Archunan, “A combinatorial model for effective estrus detection in Murrah buffalo,” Vet World, vol. 10, no. 2, pp. 209–213, Feb. 2017, doi: 10.14202/VETWORLD.2017.209-213.

[7] Srivastava, A. K., Kumaresan, A., Mohanty, T. K., & Prasad, S. (2013). Status paper on buffalo estrus biology, India. National Dairy Research Institute, Karnal, 12.

[8] S. Singha et al., “Salivary cell-free HSD17B1 and HSPA1A transcripts as potential biomarkers for estrus identification in buffaloes (Bubalus bubalis),” Anim Biotechnol, vol. 34, no. 7, 2023, doi: 10.1080/10495398.2022.2105228.

[9] G. Naidu Surla, L. K. Kumar, V. Gowdar Vedamurthy, D. Singh, and S. K. Onteru, “Salivary TIMP1 and predicted mir-141, possible transcript biomarkers for estrus in the buffalo (Bubalus bubalis),” Reprod Biol, vol. 22, no. 2, 2022, doi: 10.1016/j.repbio.2022.100641.

[10] D. SankarGanesh, R. Ramachandran, U. Suriyakalaa, A. Ramkumar, S. Achiraman, and G. Archunan, “Heat shock protein(s) may serve as estrus indicators in animals: A conceptual hypothesis,” Med Hypotheses, vol. 117, pp. 47–49, Aug. 2018, doi: 10.1016/j.mehy.2018.06.003.

[11] S. Muthukumar et al., “Exploration of salivary proteins in buffalo: An approach to find marker proteins for estrus,” FASEB Journal, vol. 28, no. 11, 2014, doi: 10.1096/fj.14-252288.

[12] L. K. Singh et al., “Comparative Proteome Profiling of Saliva Between Estrus and Non-Estrus Stages by Employing Label-Free Quantitation (LFQ) and Tandem Mass Tag (TMT)-LC-MS/MS Analysis: An Approach for Estrus Biomarker Identification in Bubalus bubalis,” Front Genet, vol. 13, 2022, doi: 10.3389/fgene.2022.867909.

[13] V. Ambros, “The functions of animal microRNAs.,” Nature, vol. 431, no. 7006, pp. 350–5, Sep. 2004, doi: 10.1038/nature02871.

[14] K. Ranganathan and V. Sivasankar, “MicroRNAs - Biology and clinical applications,” 2014. doi: 10.4103/0973-029X.140762.

[15] H. Wang, R. Peng, J. Wang, Z. Qin, and L. Xue, “Circulating microRNAs as potential cancer biomarkers: The advantage and disadvantage,” 2018. doi: 10.1186/s13148-018-0492-1.

[16] R. Chandel, R. Saxena, A. Das, and J. Kaur, “Association of rno-miR-183-96-182 cluster with diethyinitrosamine induced liver fibrosis in Wistar rats.,” J Cell Biochem, Dec. 2017, doi: 10.1002/jcb.26583.

[17] P. Sarwalia et al., “Establishment of Repertoire of Placentome-Associated MicroRNAs and Their Appearance in Blood Plasma Could Identify Early Establishment of Pregnancy in Buffalo (Bubalus bubalis),” Front Cell Dev Biol, vol. 9, 2021, doi: 10.3389/fcell.2021.673765.

[18] A. Hebbar, R. Chandel, P. Rani, S. K. Onteru, and D. Singh, “Urinary Cell-Free miR-99a-5p as a Potential Biomarker for Estrus Detection in Buffalo,” Front Vet Sci, vol. 8, p. 249, May 2021, doi: 10.3389/FVETS.2021.643910/BIBTEX.

[19] M. Joshi et al., “Detection and quantification of TIMP1 and miR-141 through RT-LAMP and TT-LAMP in serum samples during estrous cycle in buffalo,” Reprod Biol, vol. 23, no. 4, 2023, doi: 10.1016/j.repbio.2023.100820.

[20] C. Romani et al., “Stability of circulating miRNA in saliva: The influence of sample associated pre-analytical variables,” Clinica Chimica Acta, vol. 553, 2024, doi: 10.1016/j.cca.2023.117702.

[21] A. Aryani and B. Denecke, “In vitro application of ribonucleases: Comparison of the effects on mRNA and miRNA stability,” BMC Res Notes, vol. 8, no. 1, 2015, doi: 10.1186/s13104-015-1114-z.

[22] Z. Williams et al., “Comprehensive profiling of circulating microRNA via small RNA sequencing of cDNA libraries reveals biomarker potential and limitations,” Proc Natl Acad Sci U S A, vol. 110, no. 11, pp. 4255–4260, Mar. 2013, doi: 10.1073/PNAS.1214046110.

[23] S. D. Sontakke, B. T. Mohammed, A. S. McNeilly, and F. X. Donadeu, “Characterization of microRNAs differentially expressed during bovine follicle development,” Reproduction, vol. 148, no. 3, pp. 271–283, Sep. 2014, doi: 10.1530/REP-14-0140.

[24] A. Jerome, S. M. K. Thirumaran, and S. N. Kala, “Identification of microRNAs in corpus luteum of pregnancy in buffalo (Bubalus bubalis) by deep sequencing.,” Iran J Vet Res, 2017, doi: 10.22099/ijvr.2017.4637.

[25] P. Singh et al., “Salivary miR-16, miR-191 and miR-223: intuitive indicators of dominant ovarian follicles in buffaloes,” Molecular Genetics and Genomics, vol. 292, no. 5, pp. 935–953, Oct. 2017, doi: 10.1007/S00438-017-1323-3.

[26] J. Ioannidis and F. X. Donadeu, “Circulating microRNA Profiles during the Bovine Oestrous Cycle,” PLoS One, vol. 11, no. 6, p. e0158160, Jun. 2016, doi: 10.1371/JOURNAL.PONE.0158160.

[27] J. M. Yoshizawa, C. A. Schafer, J. J. Schafer, J. J. Farrell, B. J. Paster, and D. T. W. Wong, “Salivary biomarkers: Toward future clinical and diagnostic utilities,” 2013. doi: 10.1128/CMR.00021-13.

[28] H. Ahsan, “Biomolecules and biomarkers in oral cavity: bioassays and immunopathology,” 2019. doi: 10.1080/15321819.2018.1550423.

[29] M. A. R. St. John et al., “Interleukin 6 and interleukin 8 as potential biomarkers for oral cavity and oropharyngeal squamous cell carcinoma,” Archives of Otolaryngology - Head and Neck Surgery, vol. 130, no. 8, pp. 929–935, Aug. 2004, doi: 10.1001/archotol.130.8.929.

[30] C. S. Ang, S. Binos, M. I. Knight, P. J. Moate, B. G. Cocks, and M. B. McDonagh, “Global survey of the bovine salivary proteome: Integrating multidimensional prefractionation, targeted, and glycocapture strategies,” J Proteome Res, 2011, doi: 10.1021/pr200516d.

[31] Ó. Rapado-González, B. Majem, L. Muinelo-Romay, R. López-López, and M. M. Suarez-Cunqueiro, “Cancer salivary biomarkers for tumours distant to the oral cavity,” 2016. doi: 10.3390/ijms17091531.

[32] N. Malathi, S. Mythili, and H. R. Vasanthi, “Salivary Diagnostics: A Brief Review,” ISRN Dent, 2014, doi: 10.1155/2014/158786.

[33] R. Ravinder et al., “Saliva ferning, an unorthodox estrus detection method in water buffaloes (Bubalus bubalis),” Theriogenology, vol. 86, no. 5, pp. 1147–1155, Sep. 2016, doi: 10.1016/J.THERIOGENOLOGY.2016.04.004.

[34] P. Potla, S. A. Ali, and M. Kapoor, “A bioinformatics approach to microRNA-sequencing analysis,” Osteoarthr Cartil Open, vol. 3, no. 1, 2021, doi: 10.1016/j.ocarto.2020.100131.

[35] A. J. Enright, B. John, U. Gaul, T. Tuschl, C. Sander, and D. S. Marks, “MicroRNA targets in Drosophila.,” Genome Biol, vol. 5, no. 1, 2003, doi: 10.1186/gb-2003-5-1-r1.

[36] L. Chang, G. Zhou, O. Soufan, and J. Xia, “miRNet 2.0: Network-based visual analytics for miRNA functional analysis and systems biology,” Nucleic Acids Res, vol. 48, no. W1, 2020, doi: 10.1093/nar/gkaa467.

[37] J. H. Bahn et al., “The landscape of MicroRNA, piwi-interacting RNA, and circular RNA in human saliva,” Clin Chem, vol. 61, no. 1, pp. 221–230, Jan. 2015, doi: 10.1373/CLINCHEM.2014.230433.

[38] K. L. Burgos et al., “Identification of extracellular miRNA in human cerebrospinal fluid by next-generation sequencing,” RNA, vol. 19, no. 5, pp. 712–722, May 2013, doi: 10.1261/RNA.036863.112.

[39] J. E. Freedman et al., “Diverse human extracellular RNAs are widely detected in human plasma,” Nat Commun, vol. 7, Apr. 2016, doi: 10.1038/NCOMMS11106.

[40] N. A. Hasan et al., “Microbial community profiling of human saliva using shotgun metagenomic sequencing,” PLoS One, vol. 9, no. 5, 2014, doi: 10.1371/journal.pone.0097699.

[41] N. Spielmann et al., “The human salivary RNA transcriptome revealed by massively parallel sequencing,” Clin Chem, vol. 58, no. 9, 2012, doi: 10.1373/clinchem.2011.176941.

[42] A. Yeri et al., “Total extracellular small RNA profiles from plasma, saliva, and urine of healthy subjects,” Sci Rep, vol. 7, 2017, doi: 10.1038/srep44061.

[43] P. Chugh and D. P. Dittmer, “Potential pitfalls in microRNA profiling,” 2012. doi: 10.1002/wrna.1120.

[44] L. Valihrach, P. Androvic, and M. Kubista, “Circulating miRNA analysis for cancer diagnostics and therapy,” 2020. doi: 10.1016/j.mam.2019.10.002.

[45] K. E. Kaczor-Urbanowicz et al., “Novel approaches for bioinformatic analysis of salivary RNA sequencing data for development,” Bioinformatics, vol. 34, no. 1, 2018, doi: 10.1093/bioinformatics/btx504.

[46] S. Gebremedhn et al., “MicroRNA Expression Profile in Bovine Granulosa Cells of Preovulatory Dominant and Subordinate Follicles during the Late Follicular Phase of the Estrous Cycle.,” PLoS One, vol. 10, no. 5, p. e0125912, 2015, doi: 10.1371/journal.pone.0125912.

[47] J. C. da Silveira, D. N. R. Veeramachaneni, Q. A. Winger, E. M. Carnevale, and G. J. Bouma, “Cell-secreted vesicles in equine ovarian follicular fluid contain mirnas and proteins: A possible new form of cell communication within the ovarian follicle,” Biol Reprod, vol. 86, no. 3, 2012, doi: 10.1095/biolreprod.111.093252.

[48] S. Ponsuksili et al., “Differential expression of miRNAs and their target mRNAs in endometria prior to maternal recognition of pregnancy associates with endometrial receptivity for in vivo- and in vitro-produced bovine embryos,” Biol Reprod, vol. 91, no. 6, 2014, doi: 10.1095/biolreprod.114.121392.

[49] A. Lange-Consiglio et al., “Oviductal microvesicles and their effect on in vitro maturation of canine oocytes,” Reproduction, vol. 154, no. 2, pp. 167–180, 2017, doi: 10.1530/REP-17-0117.

[50] J. Zhou et al., “The let-7g microRNA promotes follicular granulosa cell apoptosis by targeting transforming growth factor-β type 1 receptor,” Mol Cell Endocrinol, vol. 409, 2015, doi: 10.1016/j.mce.2015.03.012.

[51] R. Cao, W. J. Wu, X. L. Zhou, P. Xiao, Y. Wang, and H. L. Liu, “Expression and preliminary functional profiling of the let-7 family during porcine ovary follicle Atresia,” Mol Cells, vol. 38, no. 4, 2015, doi: 10.14348/molcells.2015.2122.

[52] D. Andrei et al., “Differential miRNA Expression Profiles in Cumulus and Mural Granulosa Cells from Human Pre-ovulatory Follicles,” MicroRNA, vol. 8, no. 1, 2018, doi: 10.2174/2211536607666180912152618.

[53] F. X. Donadeu, S. N. Schauer, and S. D. Sontakke, “Involvement of miRNAs in ovarian follicular and luteal development,” Journal of Endocrinology, vol. 215, no. 3, pp. 323–334, Dec. 2012, doi: 10.1530/JOE-12-0252.

[54] A. M. M. T. Reza et al., “Roles of microRNAs in mammalian reproduction: from the commitment of germ cells to peri-implantation embryos,” Biological Reviews, vol. 94, no. 2, 2019, doi: 10.1111/brv.12459.

[55] M. Otsuka et al., “Impaired microRNA processing causes corpus luteum insufficiency and infertility in mice,” Journal of Clinical Investigation, vol. 118, no. 5, pp. 1944–1954, May 2008, doi: 10.1172/JCI33680.

[56] X. D. Zi, J. Y. Lu, and L. Ma, “Identification and comparative analysis of the ovarian microRNAs of prolific and non-prolific goats during the follicular phase using high-throughput sequencing,” Sci Rep, 2017, doi: 10.1038/s41598-017-02225-x.

[57] N. Wu et al., “Expressed microRNA associated with high rate of egg production in chicken ovarian follicles,” Anim Genet, vol. 48, no. 2, 2017, doi: 10.1111/age.12516.

[58] X. Zou et al., “Comprehensive analysis of mRNAs and miRNAs in the ovarian follicles of uniparous and multiple goats at estrus phase,” BMC Genomics, 2020, doi: 10.1186/s12864-020-6671-4.

[59] H. B. Suo, K. C. Zhang, and J. Zhao, “MiR-200a promotes cell invasion and migration of ovarian carcinoma by targeting PTEN,” Eur Rev Med Pharmacol Sci, vol. 22, no. 13, 2018, doi: 10.26355/eurrev_201807_15398.

[60] L. Záveský et al., “Ascites-Derived Extracellular microRNAs as Potential Biomarkers for Ovarian Cancer,” Reproductive Sciences, vol. 26, no. 4, 2019, doi: 10.1177/1933719118776808.

[61] Y. Wang et al., “FOXD1 is targeted by miR-30a-5p and miR-200a-5p and suppresses the proliferation of human ovarian carcinoma cells by promoting p21 expression in a p53-independent manner,” Int J Oncol, vol. 52, no. 6, 2018, doi: 10.3892/ijo.2018.4359.

[62] H. Wang et al., “Genome -wide transcriptome profiling in ovaries of small-tail Han sheep during the follicular and luteal phases of the oestrous cycle,” Anim Reprod Sci, vol. 197, 2018, doi: 10.1016/j.anireprosci.2018.08.031.

[63] H. Yang et al., “Identification and profiling of microRNAs from ovary of estrous Kazakh sheep induced by nutritional status in the anestrous season,” Anim Reprod Sci, vol. 175, 2016, doi: 10.1016/j.anireprosci.2016.10.004.

[64] M. Zhai, Y. Xie, H. Liang, X. Lei, and Z. Zhao, “Comparative profiling of differentially expressed microRNAs in estrous ovaries of Kazakh sheep in different seasons,” Gene, vol. 664, 2018, doi: 10.1016/j.gene.2018.04.025.

[65] X. D. Zhang et al., “Characterization and differential expression of microRNAs in the ovaries of pregnant and non-pregnant goats (Capra hircus),” BMC Genomics, vol. 14, no. 1, Mar. 2013, doi: 10.1186/1471-2164-14-157.

[66] M. Hossain et al., “Identification and characterization of miRNAs expressed in the bovine ovary,” BMC Genomics, vol. 10, p. 443, Sep. 2009, doi: 10.1186/1471-2164-10-443.

[67] X. Chen, H. Chen, S. Jiang, H. Shen, C. Li, and X. Zeng, “Expression profile analysis of microRNAs during the oestrous cycle of Qira black sheep,” Thai Journal of Veterinary Medicine, vol. 51, no. 3, 2021, doi: 10.14456/tjvm.2021.56.

[68] P. Landgraf et al., “A Mammalian microRNA Expression Atlas Based on Small RNA Library Sequencing,” Cell, 2007, doi: 10.1016/j.cell.2007.04.040.

[69] T. Mishima et al., “MicroRNA (miRNA) cloning analysis reveals sex differences in miRNA expression profiles between adult mouse testis and ovary,” Reproduction, vol. 136, no. 6, pp. 811–822, Dec. 2008, doi: 10.1530/REP-08-0349.

[70] D. McBride et al., “Identification of miRNAs associated with the follicular-luteal transition in the ruminant ovary,” Reproduction, vol. 144, no. 2, pp. 221–233, Aug. 2012, doi: 10.1530/REP-12-0025.

[71] D. Salilew-Wondim et al., “The expression pattern of microRNAs in granulosa cells of subordinate and dominant follicles during the early luteal phase of the bovine estrous cycle,” PLoS One, vol. 9, no. 9, Sep. 2014, doi: 10.1371/JOURNAL.PONE.0106795.

[72] R. E. Dickinson, M. Myers, and W. C. Duncan, “Novel regulated expression of the SLIT/ROBO pathway in the ovary: Possible role during luteolysis in women,” Endocrinology, vol. 149, no. 10, 2008, doi: 10.1210/en.2008-0204.

[73] J. Uma, P. Muraly, S. Verma-Kumar, and R. Medhamurthy, “Determination of onset of apoptosis in granulosa cells of the preovulatory follicles in the bonnet monkey (Macaca radiata): Correlation with mitogen-activated protein kinase activities,” Biol Reprod, vol. 69, no. 4, 2003, doi: 10.1095/biolreprod.103.017897.

[74] M. Shiota et al., “Correlation of mitogen-activated protein kinase activities with cell survival and apoptosis in porcine granulosa cells,” Zoolog Sci, vol. 20, no. 2, 2003, doi: 10.2108/zsj.20.193.

[75] E. T. Maizels et al., “Developmental regulation of mitogen-activated protein kinase-activated kinases-2 and -3 (MAPKAPK-2/-3) in vivo during corpus luteum formation in the rat,” Molecular Endocrinology, vol. 15, no. 5, 2001, doi: 10.1210/mend.15.5.0634.

[76] S. H. Lee and S. Lee, “Change of Ras and its guanosine triphosphatases (GTPases) during development and regression in bovine corpus luteum,” Theriogenology, vol. 144, 2020, doi: 10.1016/j.theriogenology.2019.12.014.

[77] Q. Wei et al., “Thyroid hormones alter estrous cyclicity and antioxidative status in the ovaries of rats,” Animal Science Journal, vol. 89, no. 3, pp. 513–526, Mar. 2018, doi: 10.1111/asj.12950.

[78] N. N. S. B. N. M. Kamal and W. N. S. Shahidan, “Non-exosomal and exosomal circulatory MicroRNAs: Which are more valid as biomarkers?,” 2020. doi: 10.3389/fphar.2019.01500.

[79] J. Baran-Gale et al., “Addressing bias in small RNA library preparation for sequencing: A new protocol recovers microRNAs that evade capture by current methods,” Front Genet, 2015, doi: 10.3389/fgene.2015.00352.

[80] O. Bryzgunova, M. Konoshenko, I. Zaporozhchenko, A. Yakovlev, and P. Laktionov, “Isolation of cell-free mirna from biological fluids: Influencing factors and methods,” 2021. doi: 10.3390/diagnostics11050865.

[81] A. D. Jayaprakash, O. Jabado, B. D. Brown, and R. Sachidanandam, “Identification and remediation of biases in the activity of RNA ligases in small-RNA deep sequencing,” Nucleic Acids Res, vol. 39, no. 21, 2011, doi: 10.1093/nar/gkr693.

[82] Y. Fu, P. H. Wu, T. Beane, P. D. Zamore, and Z. Weng, “Elimination of PCR duplicates in RNA-seq and small RNA-seq using unique molecular identifiers,” BMC Genomics, vol. 19, no. 1, 2018, doi: 10.1186/s12864-018-4933-1.

[83] C. Wright et al., “Comprehensive assessment of multiple biases in small RNA sequencing reveals significant differences in the performance of widely used methods,” BMC Genomics, vol. 20, no. 1, 2019, doi: 10.1186/s12864-019-5870-3.

[84] C. A. Raabe, T. H. Tang, J. Brosius, and T. S. Rozhdestvensky, “Biases in small RNA deep sequencing data,” Nucleic Acids Res, vol. 42, no. 3, 2014, doi: 10.1093/nar/gkt1021.

[85] N. Kosaka, H. Iguchi, and T. Ochiya, “Circulating microRNA in body fluid: a new potential biomarker for cancer diagnosis and prognosis,” Cancer Sci, vol. 101, no. 10, pp. 2087–2092, 2010, doi: 10.1111/j.1349-7006.2010.01650.x.

[86] L. Moldovan, K. E. Batte, J. Trgovcich, J. Wisler, C. B. Marsh, and M. Piper, “Methodological challenges in utilizing miRNAs as circulating biomarkers,” J Cell Mol Med, vol. 18, no. 3, 2014, doi: 10.1111/jcmm.12236.

[87] E. Presser, M. Simuyandi, and J. Brown, “The effects of storage time and temperature on recovery of salivary secretory immunoglobulin A,” American Journal of Human Biology, vol. 26, no. 3, 2014, doi: 10.1002/ajhb.22525.

